# Retinal glial cells under hypoxia possibly function in mammalian myopia by responding to mechanical stimuli, affecting hormone metabolism and peptide secretion

**DOI:** 10.1101/2020.11.26.399519

**Authors:** Xuhong Zhang, Yingying Wen, Le Jin, Dongyan Zhang, Liyue Zhang, Chen Xie, Dongyu Guo, Jianping Tong, Ye Shen

**Affiliations:** Ophthalmology department, the First Affiliated Hospital of Zhejiang University, Hangzhou, ZhejiangProvince, China

**Keywords:** Gene set enrichment analysis, Myopia, astrocyte, Müller cell

## Abstract

**Purpose:** Changes in the retina and the choroid blood vessels are regularly observed in myopia. The aim of this study is to test if the retinal glial cells, which directly contact blood vessels, play a role in mammalian myopia.

**Method:** We adapted the common form-deprivation myopia mouse model and used the retina slice and whole-mount immunofluorescence technique to evaluate changes in the morphology, and distribution of retinal glial cells. We then searched the Gene Expression Omnibus database for series on myopia and retinal glial cells (astrocytes and Müller cells). Using review articles and the National Center for Biotechnology Information gene database, we obtained clusters of myopia-related gene lists. By searching SwissTargetPrediction, we collected information on atropine target proteins. We then used online tools to find the Gene Ontology analysis enriched clusters, pathways, and proteins that provide correlative evidence.

**Result:** Glial fibrillary acidic protein fluorescence was observed in mice eyes that were covered and deprived of light for five days compared to uncovered eyes, and the cell morphology became more star-like. From in silico experiments, we identified several pathways and proteins that were common to both myopia and retinal glial cells in hypoxic conditions. The common pathways in human astrocytes represented response to mechanical stimuli, peptide secretion, skeletal system development, and monosaccharide binding. In mouse Müller cells, the pathways represented membrane raft and extracellular structure organisation. The proteins common to myopia and hypoxic glial cells were highly relevant to atropine target proteins.

**Conclusion:** Retinal astrocytes and Müller cells under hypoxic conditions contribute to the development of myopia, and may be a valid target for atropine.

## Introduction

Myopia, which can cause severe fundus oculi diseases and even blindness^[1]^, is becoming a global problem^[2]^. Numerous studies have uncovered many aspects and underlying mechanisms of myopia^[3]^. Scleral expansion whereby the ocular axis becomes longer is commonly considered a consequence of myopia^[4]^. During the process, retinal and choroid vessel changes have been observed where not only the density is altered, but also the blood flow and oxygen content^[5–7]^. The decrease in vessel density and oxygen content cause the ocular hypoxia condition. Retinal glial cells are widely distributed and have many functions^[8]^. Astrocytes are in close contact with blood vessels, and Müller cells provide radial support. Retinal Müller cells and astrocytes have also been abundantly tested in a variety of ocular diseases, including retinal detachment, light damage, and ischemia^[9, 10]^, in which they show increased glial fibrillary acidic protein (GFAP) expression and morphological changes. Activated astrocytes can produce nitric oxide, tumour necrosis factor-alpha (TNF-α), reactive oxygen species, and other neurotoxic mediators^[11]^. There are few reports of the relationship between retinal glial cells and myopia, although several pathological process, including oxidative stress^[12]^ and hypoxia^[13]^, are associated with myopia.

Atropine is one of the most effective drugs used to delay myopia progression. The muscarinic receptors (MRs) and several non-MRs^[14]^ found mainly in the retina and sclera have been proposed as targets of atropine. In malathion-induced brain injury, atropine decreased brain astrocyte GFAP expression by inhibiting nitric oxide rather than inhibiting brain acetylcholine esterase activity^[11]^. We cannot exclude that atropine may benefit myopia by targeting the glial cells. Therefore, we hypothesised that retinal glial cells may have a role in myopia and may be involved in atropine intervention. To test this hypothesis, we used the form-deprivation myopia (FDM) mouse model and investigated retinal GFAP expression. Furthermore, we explored high-throughput data on myopia and retinal glial cell performance under hypoxic conditions to analyse their function and potential mechanisms involved.

## Materials & Methods

To test our hypothesis in vivo and in silico, we used the FDM mouse model and searched many databases to obtain sufficient relevant data. Figure 1 presents the experimental flowchart.

**Figure 1:**
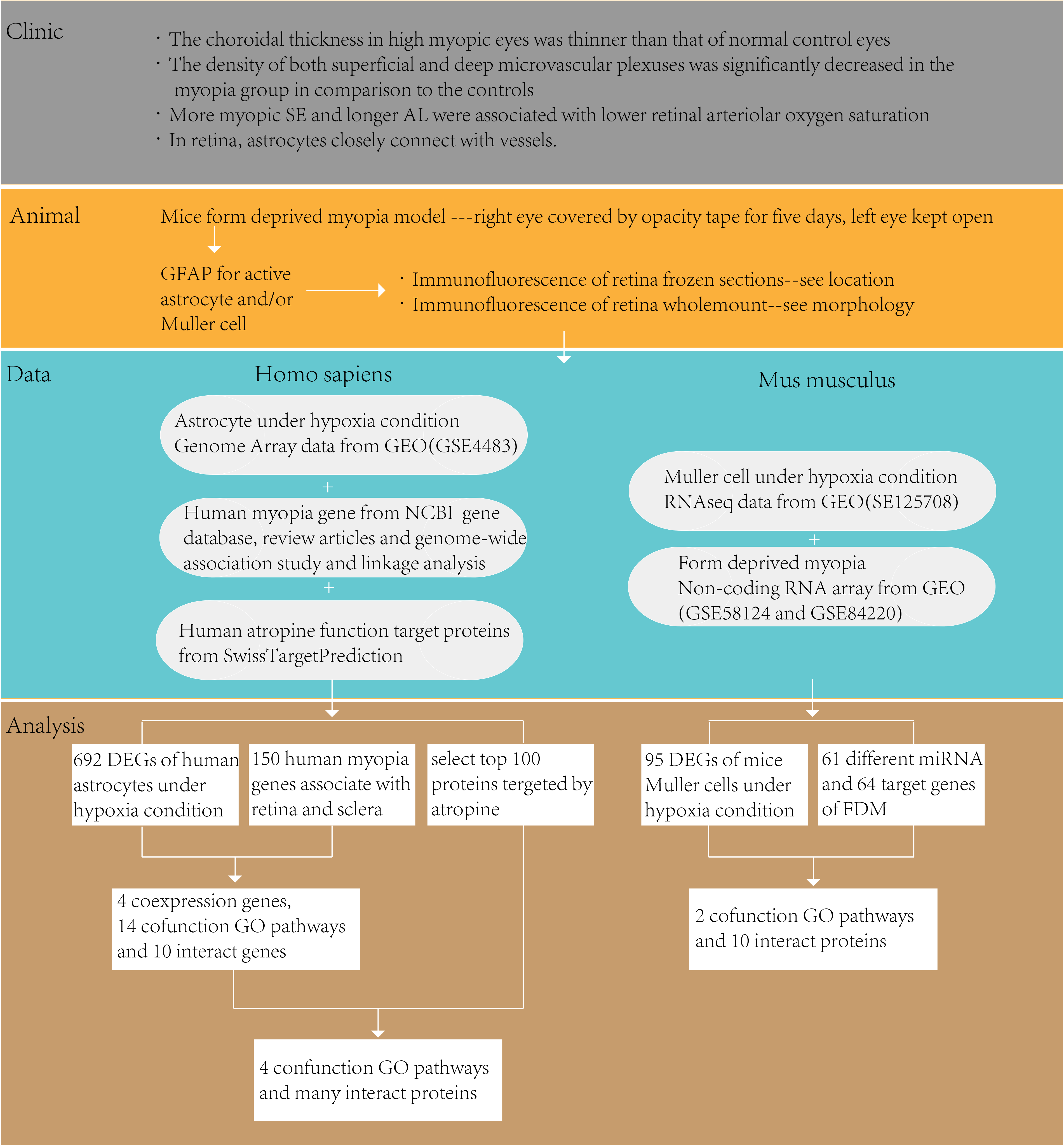
Flowchart of the experimental process.

### Mouse form-deprivation myopia

We followed the widely used method to induce FDM in C57BL/6 mice^[15]^. Postnatal day 20 mice were raised in a 12/12 light/dark cycle and room temperature of 26°C. Food and water were provided ad libitum. The right eyes of mice were covered with opaque packing tape for five days and the left eyes were left uncovered. The mice were checked daily to make sure the covers had not come off. All procedures were carried out in accordance with the National Institutes of Health Guidelines for the Care and Use of Laboratory Animals and were approved by the Animal Advisory Committee at Zhejiang University (Approval No. ZJU20200118).

### Immunofluorescence staining in retina frozen sections and whole-mounts

After five days, the mice were anesthetised with pentobarbitone (50 mg/kg intraperitoneal). When the mice no longer responded to hard nipping of their tails, myocardial perfusion was performed and the eyeballs were removed and fixed in 4% (w/v) paraformaldehyde in phosphate-buffered saline (PBS) for two hours at 4°C. The anterior part of the ocular, including the conea, iris and lens, was dissected under a microscope, and the optic cup was suspended in 20–30% (w/v) sucrose solution. Once the cup settled to the bottom, it was transferred to a higher concentration of sucrose solution. After dehydration, the optic cups were positioned in embedding medium (Neg-50; Thermo Scientific, Waltham, MA, USA) and frozen. We collected 15-μm-thick cryosections and air dried the slides at room temperature, after which they were washed with PBS for 10 min, dried, and blocked with confining liquid (10% normal donkey serum (NDS), 1% bovine serum albumin (BSA), and 0.3% triton X-100 in PBS) for one hour at room temperature in a humidity chamber. The sections were then dried, and the primary antibody anti-GFAP (16825-1-AP; Proteintech, Subsidiary in China) was diluted 1:300 with antibody diluent (1% NDS, 1% BSA, and 0.3% triton in PBS) and added onto the slides, which were incubated for one hour at room temperature. The slides were washed with PBS three times (five min each time), and then incubated with Alexa Fluor 488-conjugated secondary antibody Donkey anti-Rabbit IgG (1:1000; A-21206; Thermo Fisher Scientific) for one hour at room temperature. After washing in PBS and drying, DAPI (1:4000; C1002; Beyotime, Shanghai, China) was used to label the nuclei. The sections were then rinsed three times with PBS, and mounted with 60% glycerinum under coverslips. For retina whole-mount, retina tissues didn’t need dehydration, the staining procedure was the same. Fluorescence images were acquired using a confocal microscope (FV1000; Olympus, Tokyo, Japan). The quantitation of the images was performed using the ImageJ software. Data statistic analysis was performed with SPSS (IBM SPSS statistic 21), and p<0.05 was set as significant different. Panals were put into multi-part figures with Adobe Illustrator CC 2018.

### Data search

We primarily focused on the Gene Expression Omnibus (GEO) database and searched updated series for “myopia”, “astrocyte”, and “Müller”. After manual selection, we found four available series: GSE58124 and GSE84220 for mouse FDM, GSE125708 for the mouse chronic hypoxia Müller cell gene, and GSE4483 for human astrocyte function under hypoxia. For human myopia genes, we searched the National Center for Biotechnology Information (NCBI) gene database (https://www.ncbi.nlm.nih.gov/gene/?term=myopia), genome-wide association studies and linkage analyses^[16, 17]^, as well as review articles. We also collected atropine functional target proteins using SwissTargetPrediction (http://new.swisstargetprediction.ch/result.php?job=477387596&organism=Homo_sapiens).

### Bioinformatics analysis

After finding the proper series, we downloaded the matrix files, or the results of online GEO2R analysis with the information file of the sequence platform. We then used R (version 3.6.3) for pre-processing. For the differential gene analyses, we identified differentially expressed genes (DEGs) based on a p-value less than 0.05 and fold-change (absolute value of logFC) equal to or more than 2. To find the trend of total expression of the human myopia-related gene (HMRG) in astrocytes under hypoxia, we applied the analysis principle of Gene Set Enrichment Analysis (GSEA) to make an HMRG enrichment plot. We took the total 150 HMRGs as a reference group (Gene Ontology [GO] group), and found the astrocyte gene distribution and trend in this group. From GSE58124 and GSE84220, we found 61 differentially expressed miRNAs. miRNA does not usually encode proteins, so we could not directly perform functional enrichment analysis. Instead, we analysed their functions through their corresponding mRNA using the online tool miRpath (http://www.microrna.gr/miRPathv3/)^[18]^, which also provides the results of target genes. With genes, we performed GO analysis using another online tool, Metascape (http://metascape.org/gp/index.html#/main/step1)^[19]^. With its additional protein–protein interaction (PPI) results, we found the specific proteins and their functions in the associated pathways. By combining functional protein clusters using a third online tool STRING (https://stringdb.org/cgi/input.pl?sessionId=UsvFU73mNPKS&input_page_show_search=on), we determined if the protein clusters interact with each other. The gene sequence GSE4483 was directly analysed using Metascape. With the genes related to myopia, hypoxic glial cells, and atropine target proteins, we analysed the contribution of atropine to myopia-related glial cells.

## Results

### Immunofluorescence of astrocytes and Müller cells from FDM mice

Immunofluorescence staining using GFAP as a specific marker for astrocytes and Müller cells, as well as DAPI for nuclei, was conducted for covered and exposed eyes of FDM mice. GFAP fluorescence in the FDM retina sections (Fig. 2 B) was much higher than that in the lateral open eye (Fig. 2 A). In the open-eye retina, astrocytes were on the inner side of the retinal ganglion cells (RGCs), while in the covered eye, the horizontal area was larger and displayed some thin fibres deep into the inner plexiform layer (IPL), which was much thicker than that of the open eye. In those fibres, there was DAPI fluorescence in soma-like structures, suggesting either migration or expansion of astrocytes. In whole-mount retina, we also observed GFAP enrichment in astrocytes in the FDM eye. The morphology was more star-like, and their connection with retinal vessels was obvious. However, in the open eye, GFAP was not as clearly visible and cell morphology was not distinct (Fig. 2 C and D). From the zoom images (88×), there was no significant enlargement of the domain area, but there was more overlap in that area (Fig. 2 E and F). Statistical analysis of fluorescence intensity in FDM and open eyes showed a significant difference, after FDM, the GFAP area enlarged (p=0.0015, n=6) (Fig. 2 G), and the mean fluorescence intensity increased (p = 0.075, n = 6) (Fig. 2 H).

**Figure 2:**
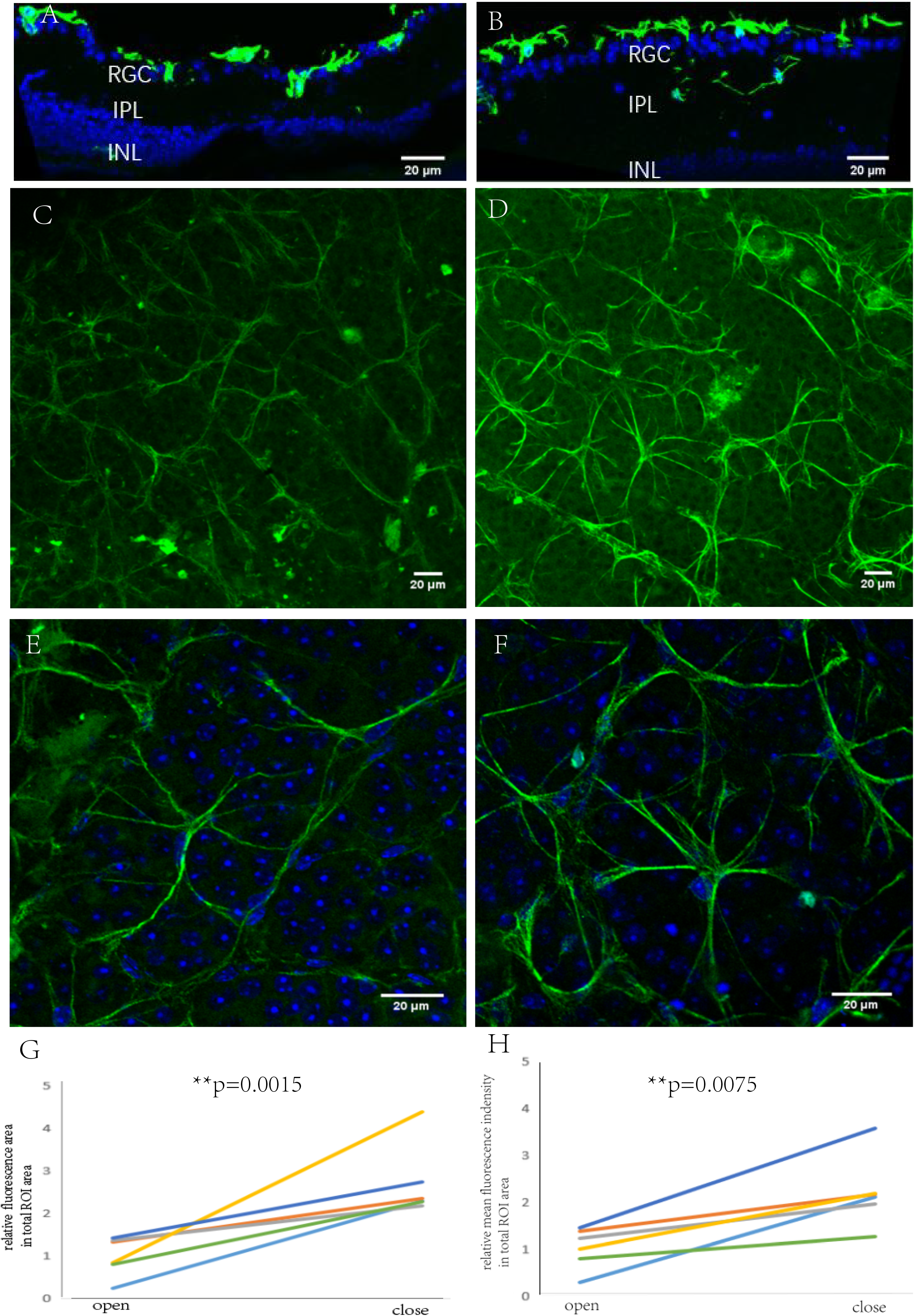
Immunofluorescence of GFAP in FDM mice. (A) section of the open eye, (B) section of the FDM eye, (C) whole-mount of the open eye, (D) whole-mount of the FDM eye, (E) zoom image of open eye whole-mount, (F) zoom image of FDM eye whole-mount, (G) percentage of area of GFAP fluorescence, (H) analysis of section mean fluorescence intensity of six mice.

### Myopia-related genes and hypoxic astrocyte-related genes in human

Human myopia-related genes have been studied widely. A total of 217 human myopia genes are listed in the NCBI gene database, and 172 genes are listed in the NHGRI GWAS Catalogue and ClinVar. The genes were systematically grouped by Tedja et al.^[20]^ according to expression position. Because our research focused on the retina and sclera, we selected all retina-related genes together with 27 pathologic myopia-related genes listed by Wu et al.^[13]^. We took hypoxic astrocyte-related DEGs from the GEO database (GSE4483). In total, we obtained 692 genes for hypoxic astrocytes and 150 genes for human myopia.

Four genes were shared between the two groups of the Venn diagram (Fig. 3 A): *GRIA4, RP2, CNGB3*, and *ADAMTS10*. *GRIA4* is expressed in cone ON bipolar cells and is responsible for common refractive error. *RP2* and *CNGB3* are expressed in cones and rods, and are associated with syndromic myopia. *ADAMTS10* is expressed in the sclera. Furthermore, expression of all these genes increased when astrocytes suffered from hypoxia. As shown in Figure 3 B, although the two groups covered all genes, the interaction between the groups was not uniform. By applying the analysis principle of the GSEA, we found that most HMRG enrichment was in the non-hypoxic part, indicating that most HMRG expression decreased when astrocytes suffered from hypoxia (Fig. 3 C).

**Figure 3.**
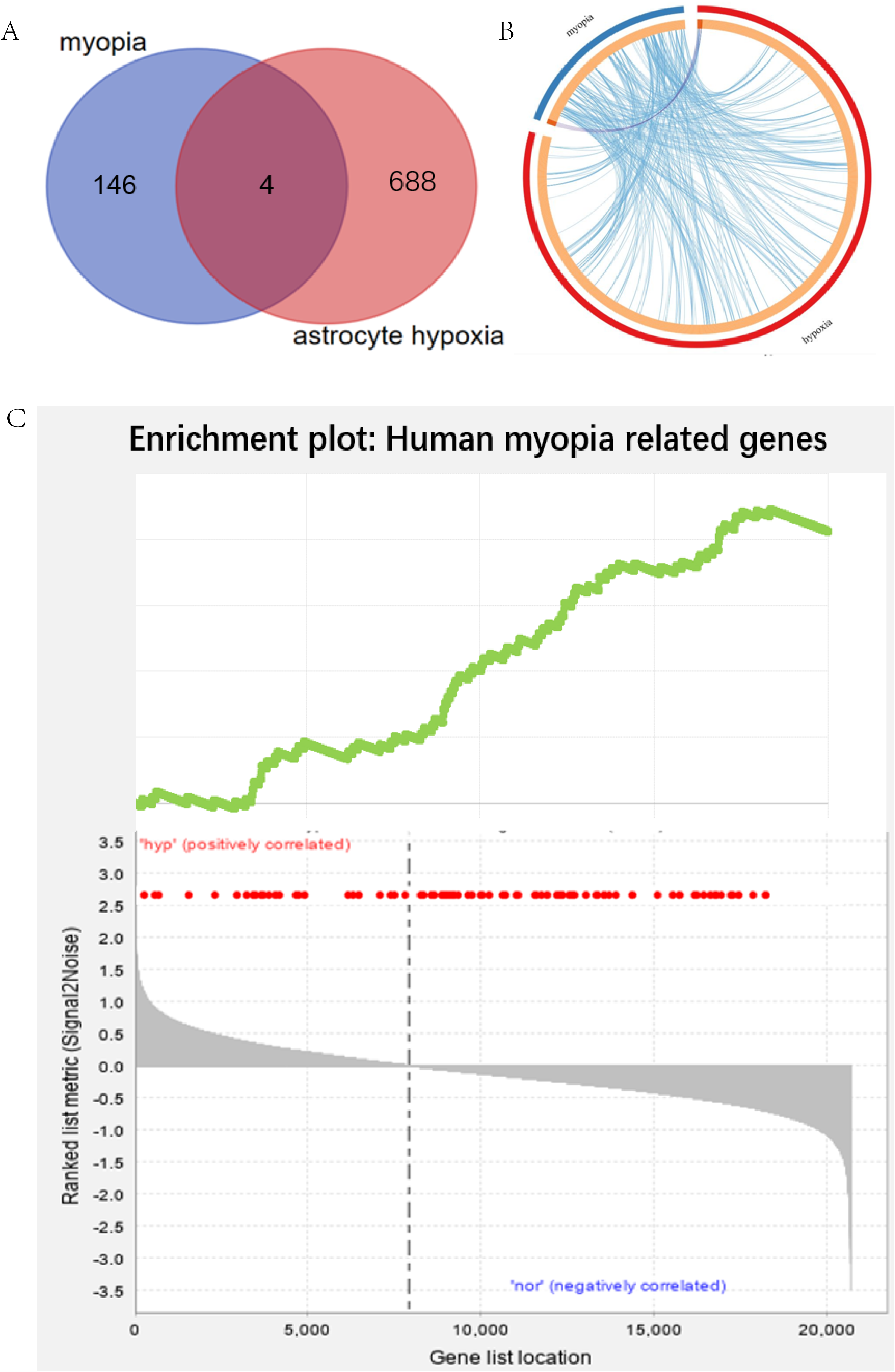
Sum up of DEGs in human myopia and astrocyte hypoxia. (A) Venn diagram and (B) chordal graph as well as GSEA analysis (C) shows the distribution and expression trend of myopia and hypoxia astrocyte genes.

The PPI analysis revealed a strong connection of the HMRGs (red nodes) with hypoxic astrocyte-related genes (Fig. 4). Because most of the myopia genes we selected were confirmed by other researchers, the connections we observed were more valid. Genes expressed in astrocytes, such as *ADM, ACTM2, FECH, EFNA3, EFNA5, MYB, MAF, ERBB2, RORA*, and *PPARGC1A*, also showed strong connections with HMRGs.

**Figure 4.**
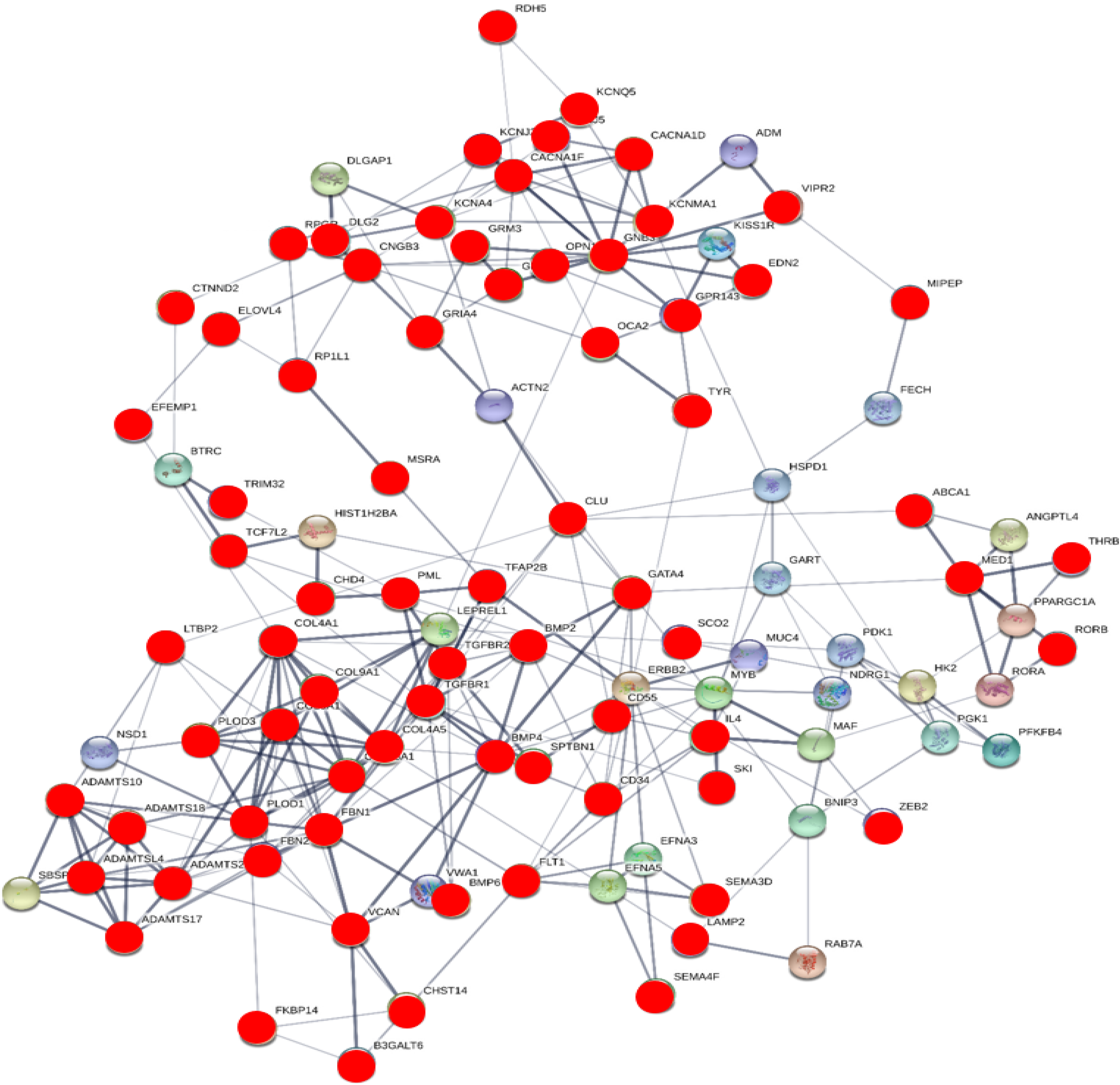
Partial PPI analysis. Red nodes represent HMRGs, and thicker lines between nodes indicate a stronger relationship.

### Gene pathway analysis in human myopia and hypoxic astrocytes

To further explore the connection between myopia and astrocytes, we derived the main GO cluster pathways by analysing the genes of the two groups (Fig. 5). The top 12 identified genes were involved in response to mechanical stimuli, peptide secretion, pigment metabolism, cellular and hormone metabolic processes, receptor regulation activity, sulphur compound binding, angiogenesis, regulation of anatomical structure and size, apoptotic signalling pathway, inorganic ion homeostasis, maintenance of protein localisation, and embryonic cranial skeleton morphogenesis.

**Figure 5.**
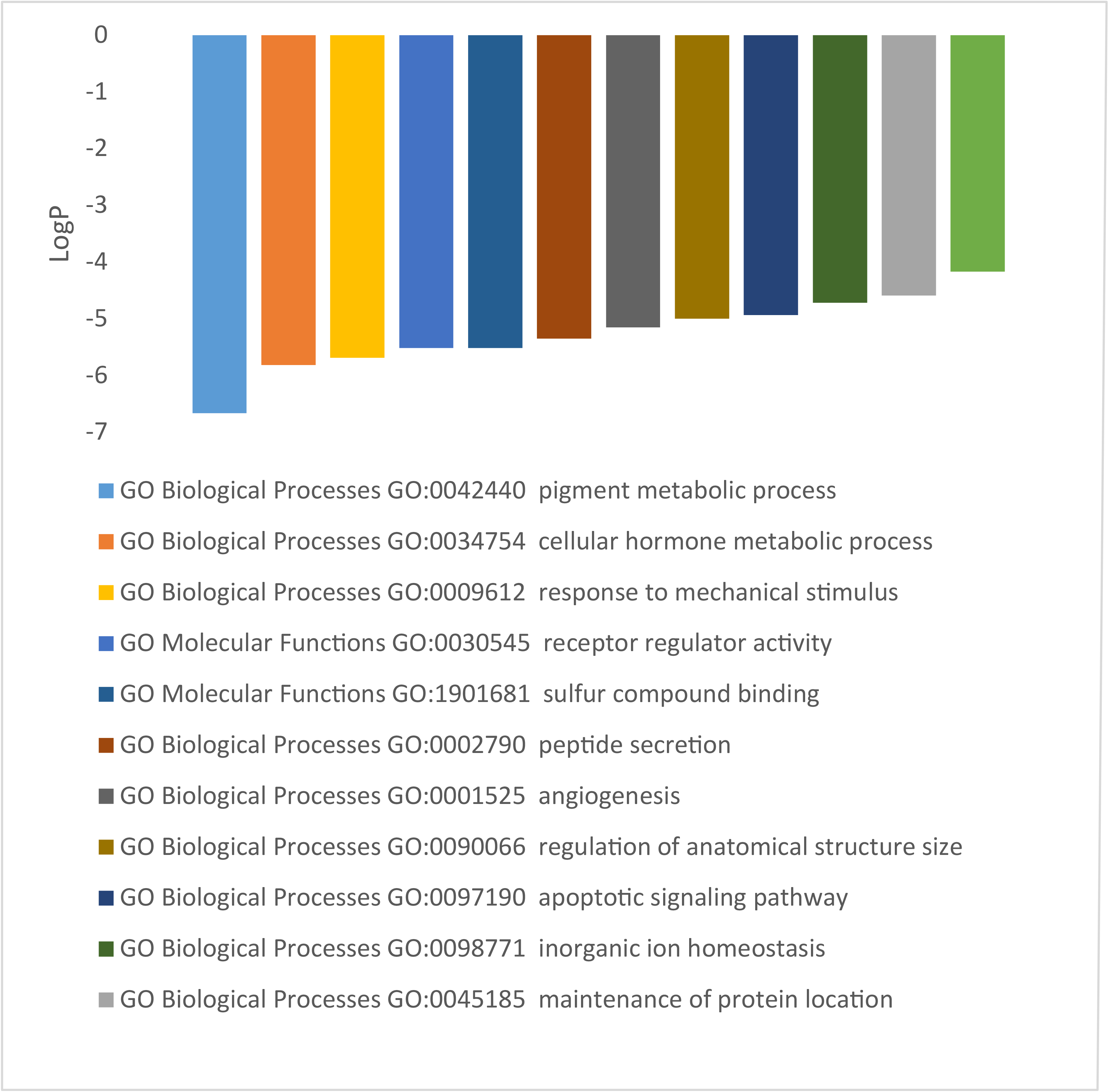
GO pathway analysis showing common pathways in the top 12 clusters.

### Atropine targets proteins with roles in myopia and astrocyte function

The different mechanisms that underlie the effectiveness of atropine against myopia are widely accepted. We collected a list of target proteins from SwissTargetPrediction and chose the top 100 as the most relevant. By comparing the gene lists for myopia, astrocytes, and atropine, we found some pathways that were common in any two of the selected lists, and some pathways even common in all three lists. Examples included pathways involved in inorganic ion homeostasis, peptide secretion, response to mechanical stimuli, and adipocyte differentiation (Fig. 6 A). Excluding myopia, atropine had four GO clusters related to astrocytes. These clusters included neuronal death, ageing, positive regulation of the MAPK cascade, and maintenance of protein localisation. The protein PPI analysis revealed the protein functions in clusters (Fig. 6 B). As there is a lot of evidence that supports the effectiveness of atropine for the treatment of myopia, this indirectly suggests that retinal glial cells may play a role in myopia.

**Figure 6.**
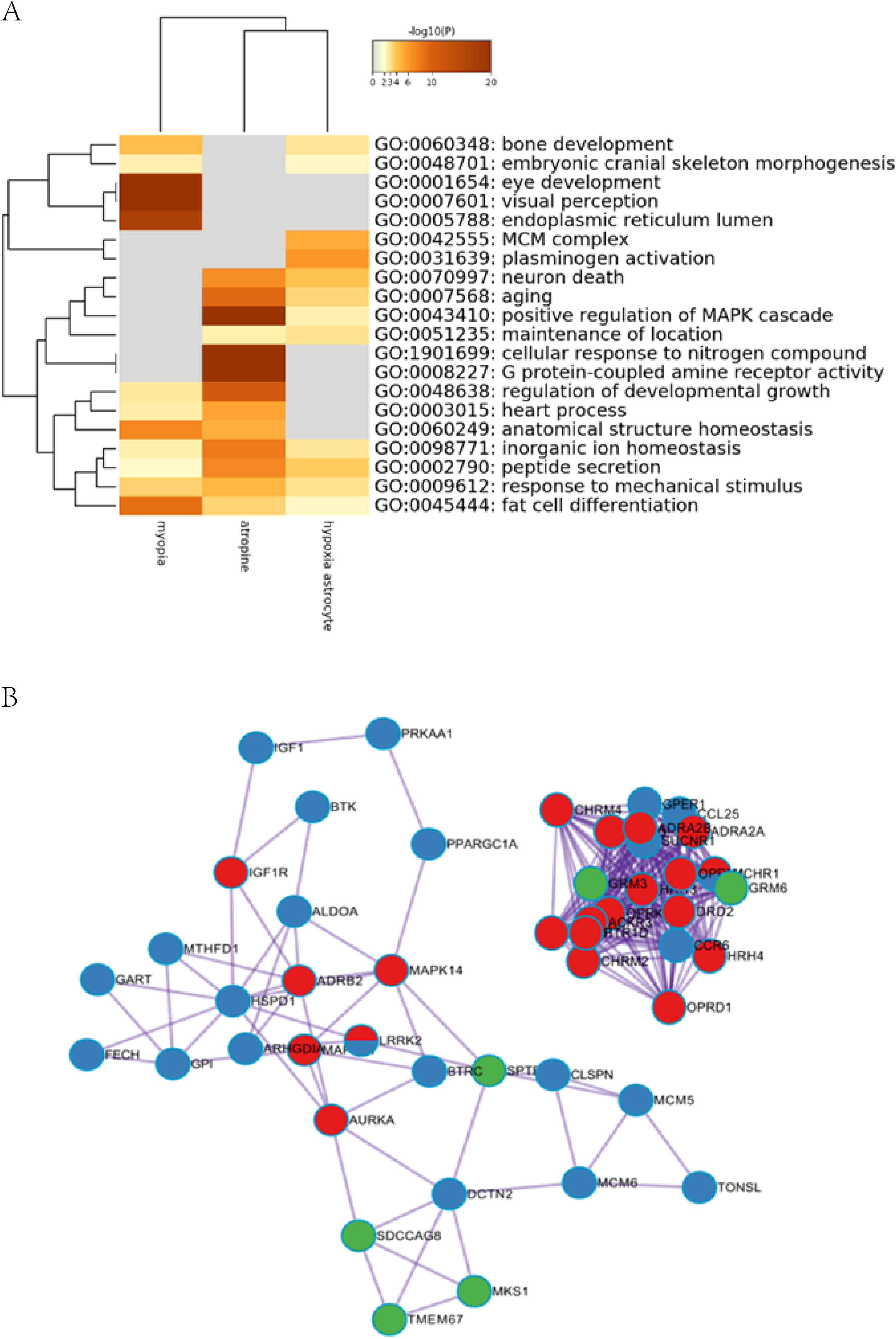
Effect of atropine on myopia and astrocytes. (A) GO analysis of the common pathways, and (B) PPI analysis of the connected nodes (red: atropine target proteins, blue: myopia-related proteins, green: astrocyte-related proteins).

### Myopia-related genes and hypoxic Müller cell-related genes in mice

We combined two similar databases, for which the primary data were miRNA expression in FDM mice. After analysis with miRpath, we generated a heat-map of 33 miRNA by Kyoto Encyclopedia of Genes and Genomes (KEGG) analysis. The main different miRNAs were mmu-miR-1187, mmu-miR-574-5p, mmu-miR-466hj, mmu-miR-325-3p, mmu-miR-465b-5p, and mmu-miR-465f-5p (Fig. 7 A). Based on the differentially expressed miRNAs, we identified the target genes in FDM mice. The five most significant pathways were GABAergic synapse, extracellular structure organisation, heterotrimeric G-protein complex, protease binding, and 3’,5’-cyclic nucleotide phosphodiesterase activity. With the supplemental materials from the GEO database (GSE125708), we identified the target genes in hypoxic Müller cells. Both myopia and hypoxia in mice have relatively centralised function genes and proteins with few interactions in between. Membrane raft and extracellular structure organisation were two GO cluster pathways common between the two groups (Fig. 7 B). The genes *Lpar1, S1pr1, Ednrb, Rgs2, Fzd5, Apln, Gapdh, Vim, Id1,* and *Vegfa* expressed by Müller cells code for proteins that link the two main functions (Fig. 7 C). They function mainly in the cytoplasm where they bind to protein and have roles in the regulation of localisation, cellular response to stimuli, and regulation of multicellular organismal processes.

**Figure 7.**
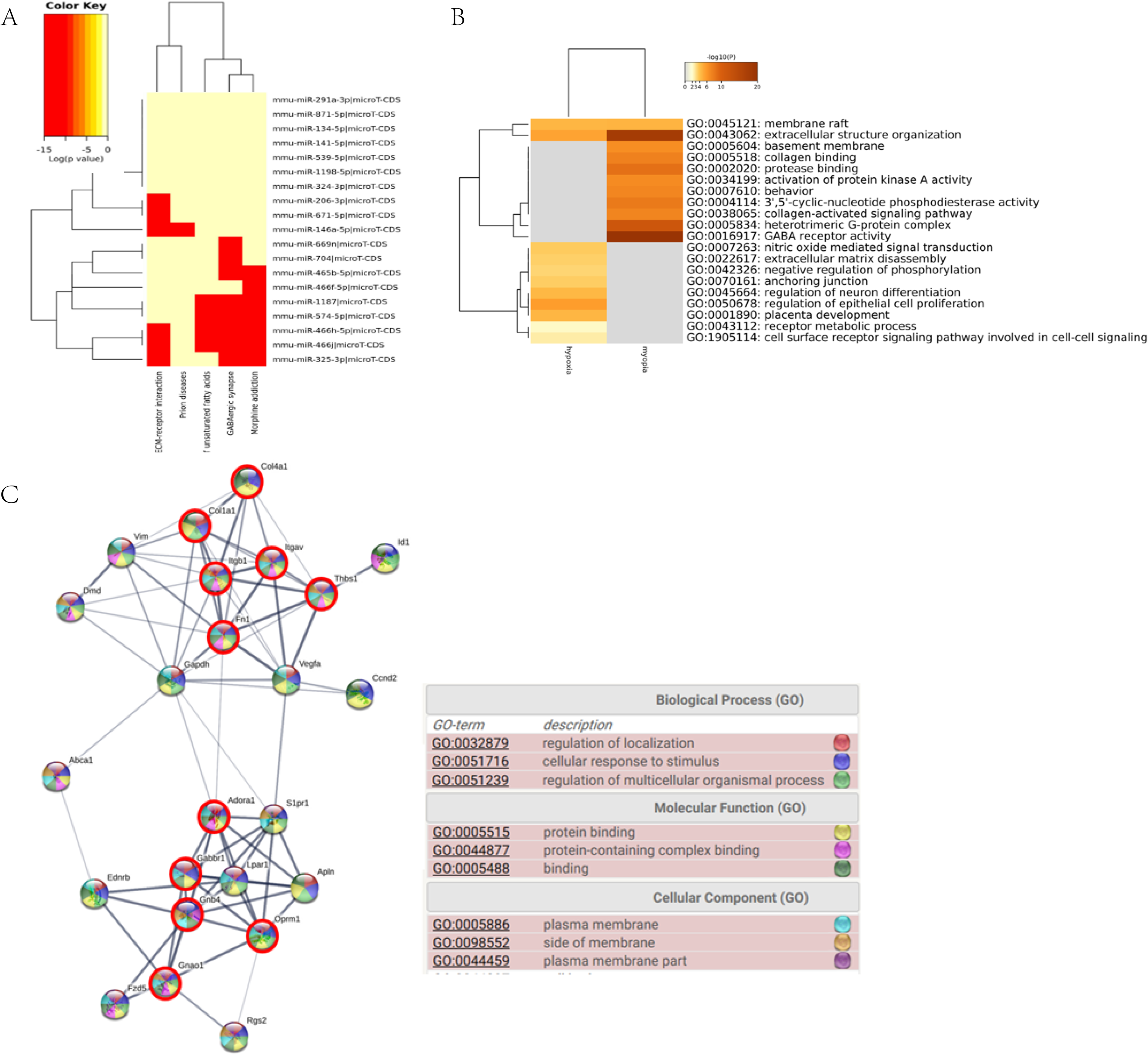
Myopia-related genes and hypoxic Müller cell-related genes in mice: (A) miRNA clusters in FDM mice, (B) the common pathways of the two conditions, and (C) PPI analysis and GO annotation results (nodes with a red circle represent myopia-related genes).

## Discussion

The main results we obtained from the in vivo and in silico experiments were that in myopia, retinal glial cells, namely astrocytes and Müller cells, play a role under hypoxic conditions. The functional pathways we identified include response to mechanical stimuli and peptide secretion.

### Hypoxia in myopia

Oxygen is transported in blood vessels, and ocular vessels have long been studied in myopia. With the development of optical coherence tomography technology, ophthalmologists and scientists can examine a larger area of ocular vessels, especially retinal and choroid vessels. The horizontal and vertical axes, as well as morphology and vascular oxygen saturation, can be examined^[5, 6, 21, 22]^. Studies have reported that choroid thickness tends to decrease in myopic patients^[7]^ and in experimentally induced myopia^[23]^. Furthermore, the density of retinal vessels is also decreased in myopia^[5]^. Although it is unascertainable that hypoxia is the cause or outcome of myopia, the two conditions are strongly connected. In our retina whole-mount GFAP photofluorogram, we observed widened blood vessel outlines by distinguishing astrocyte termini, implying vessel change. A study that focused on the sclera concluded that the activation of the scleral hypoxia pathway causes myopia, and that scleral hypoxia could be a target for myopia therapy^[13]^. From the perspective of ocular blood supply, there is a close relationship between the retina and scleral hypoxia. All ocular vessels arise from the internal carotid next to the ophthalmic artery. The two important vessels that supply the inner retina and outer retina-choroid-sclera are the central artery of the retina and the short posterior ciliary artery, respectively. Although their diameters are not large, they emerge directly from the ophthalmic artery are easily perfused; therefore, there is less risk in the case of angiemphraxis. When the retina oxygen consumption increases, the short posterior ciliary artery supplies more oxygen to the outer retina and consequently less to the sclera. Hence, part of the retina and the outer ocular layers share the same fate. Tkatchenko et al.^[24]^ found that the main pathways associated with myopia and ocular axis elongation are cell growth, and maintenance and nucleic acid metabolism. It was also reported that photoreceptor differentiation from neural progenitors occurred in the outer retina^[24]^. A study on chickens found that the main functional pathways associated with myopia were related to cell energy metabolism and synaptic transmission^[25]^. These findings regarding myopia suggest that in responding to specific visual signals, the outer retina photoreceptors are remodelled and consume more oxygen to generate energy, leading to hypoxia in the retina and the sclera. This stimulates the retinal glial cells to release visual signal dependent factors to the outer ocular layers. These factors together with the scleral factors cause final scleral remodelling. Our results also support the hypoxia function in myopia through many different pathways. In summary, ocular hypoxia may play an essential role in myopia.

### Retinal glia cells function in myopia

Research on retinal glial cells in myopia, especially astrocytes, is sparse. As mentioned previously, retinal glial cells are expected to have an important role in retina signal transduction^[26]^. For Müller cells, A study reported that after hundreds of minutes of retina stretch, Müller cells secreted c-Fos and bFGF, which are stretch-time dependent^[27]^. Stretching of the retina is thought to be a myopia risk factor during accommodation. Also, AQP4 is recognised as a regulator of fluid flow in the brain and across the retina^[28]^. In chick FDM, AQP4 was detected in the end-feet of the Müller cell membrane at the vitreous interface, nerve fibre layer, ganglion cell layer, and IPL^[29]^. Therefore, it was proposed that the role of Müller cells in myopia was to regulate the vitreous and retina fluid balance. When the fluid flow into the vitreous increased, the vitreous chamber deepened and the ocular axis became longer. Furthermore, Müller cells were found to release different factors depending on the extent of hypoxia. In mild hypoxia, Müller cells synthesised a protein factor that downregulated the expression of pigment epithelium derived factors (PEDF), or its turnover, and angiogenesis was facilitated. However, under severe (or chronic) hypoxia, PEDF exhibited neurotrophic effects^[30]^.

For astrocytes, only Lin et al.^[31]^, in a meeting abstract, reported that the loss of astrocytes co-localised with capillaries across the retina, induced myopia in juvenile marmosets, together with the decrease in capillary density. This indirectly proves that the function of astrocytes changes under hypoxic conditions. In addition to their close contact with capillaries, astrocytes have a wide range of functions in neurodevelopment of both brain and retina. They produce a wide range of factors targeting the RGCs, including IL-33^[32]^, thrombospondins and brain-derived neurotrophic factor (BDNF)^[33]^. It is possible that in most experimental myopia models, the young animals experience myopia during vision neurodevelopment, and visual acuity may also have an impact during mouse postnatal day 19–32 FDM development^[34]^, in which astrocytes may play a role. While astrocytes have many functional pathways, in our analysis, both astrocytes and Müller cells shared functional pathways with myopia. Therefore, our assumption is that retinal glial cells have an important role in myopia. Müller cells tend to be activated in response to large area stimuli such as ocular stretch and ocular fluid regulation, and thus transfer vision signals to outer ocular layers, while astrocytes tend to transfer vision signals to higher vision centres by closely influencing RGCs and other neuronal cells.

### Further reading of our in vivo lab results

Based on the immunofluorescence results, we further speculated that astrocytes and Müller cells may both be involved in myopia, but astrocytes more effectively. After five days of starting the experiment with the FDM mice, no retina attenuation was observed, nor did the number of cells decrease, which suggested that there was no severe retinal damage and confirmed that FDM is an interference of vision signal. However, the GFAP intensity in the FDM group was higher, and mainly in the RGC layer with some thin fibres protruding into the thickened IPL (Fig. 6 B). Other groups have reported that GFAP was expressed in both astrocytes and Müller cells, and that the distribution of GFAP in Müller cells was polar, and primarily in the vitreous face end-feet^[9]^. Upon stimulation (by inflammation or stretching), the GFAP signal extended from the end-feet to the outer nuclear layer^[9]^. It is possible that the stimulus used in our experiments was not strong enough. This may explain why we did not observe GFAP fluorescence beyond the RGC and IPL, and why the thin fibres in the IPL were shorter. However, from the GFAP fluorescence alone, we could not distinguish if the cells we observed were solely astrocytes, or a combination of astrocytes and Müller cell end-feet. In the whole-mount samples, we observed that the astrocyte morphology had changed (Fig. 6 D and F) without extra fluorescence, unlike the Müller cell end-feet, because there was only a small degree of overlap between the Müller cell end-feet and the astrocytes, mostly concentrated on the astrocyte soma^[9]^. This finding is in agreement with previous studies on retinal detachment in which astrocyte morphology was reported to change in the pathological state^[10]^. However, in FDM, astrocytes showed increased overlap, whereas in retinal detachment they become more “ragged”^[10]^. It is possible that the functional mechanism is different, but this needs more study. Based on our findings, we suggest that astrocytes play a more important role than Müller cells, but further research is needed.

### Further reading of our in silico results

After searching the GEO database, we obtained a similar design for mouse Müller cells and human astrocytes. Therefore, we tested the hypothesis in both species. We found that the functional pathways in human astrocytes represented response to mechanical stimuli, peptide secretion, pigment metabolic processes, cellular hormone metabolic processes, receptor regulator activity, sulphur compound binding, angiogenesis, regulation of anatomical structure size, apoptotic signalling pathway, inorganic ion homeostasis, maintenance of protein localisation, and embryonic cranial skeleton morphogenesis (Fig. 5). However, in mouse Müller cells, the pathways were involved in membrane raft and extracellular structure organisation (Fig. 7 B). A previous study reported that cell proliferation activities and synaptic plasticity supersede other activities in the myopic retina^[24]^, which is consistent with our findings, as neuronal cells are the first cells to respond to light. Following cell proliferation, the retina becomes hypoxic and the functional pathways we listed are activated to elicit a response to stretch and to regulate hormone and extracellular structure components. These actions seem to encompass the immediate response following the visual signal and neural reaction, which suggests that these pathways may be the crucial step leading to myopia, and the start of a harmful feedback loop that maintains ocular axis growth. Some of the functions have been experimentally proven, such as hormone regulation. Thyroid hormone signalling activity can decide the M/S-cone fate^[35]^. Matrix metalloproteinase 2 (MMP-2) is a widely recognised extracellular remodelling factor that is generally found in the retina and scleral tissue in myopia^[36–38]^. The majority of the core proteins in mice consisted of Lpar1, S1pr1, Ednrb, Rgs2, Fzd5, Apln, Gapdh, Vim, Id1, and Vegfa (Fig. 7 C). Other proteins were mainly associated with development and cell differentiation in humans, such as EFNA3, EFNA5, MYB, MAF, and RORA (Fig. 4). For atropine in myopia and astrocytes, the four common pathways represented inorganic ion homeostasis, peptide secretion, response to mechanical stimuli, and fat cell differentiation (Fig. 6 A). Mechanical stimuli are common in the ocular system, and have been proven to influence myopia in several studies^[38, 39]^.Atropine benefits myopia through MRs which induce mechanical change^[40]^. The same occurs in astrocytes. Whether these mechanisms actually affect myopia remains unclear and needs to be addressed based on the results of our bioinformatics analyses.

Peptide secretion is important in neuronal tissue. Many peptides, including some proglucagon-derived peptides, affect FDM. Posterior eye growth was triggered in response to glucagon, oxyntomodulin, glucagon-like peptide-1, and miniglucagon^[41]^. In Egyptian patients, a significant association of the insulin-like growth factor-1 gene rs6214, polymorphism with simple myopia, and high-grade myopia was found^[42]^. Vasoactive intestinal peptide also plays a significant role in myopia^[43–45]^. Based on in silico results, we concluded that retinal glial cells can promote myopia by responding to mechanical stimuli, affecting hormone metabolism, peptide secretion, and cell differentiation.

### Study limitation and future design

Data from the GEO database may not be optimal, although we prudently selected only four series from the whole database. To better test our hypothesis, we require designed samples and our own high-sequence data. Also, we should increase the number of mice used for the validation. For future studies, we should pay more attention to retinal glial cells in myopia and conduct further studies to distinguish the difference in function between Müller cells and astrocytes. Also, we need to investigate the detailed signalling mechanism and address other cell types that may be affected by hypoxic conditions. Finally, development of visual acuity and neurodevelopment need to be investigated. Regardless, the present study serves as a modest foundation for future research efforts.

## Conclusions

We hypothesised that retinal glial cells play a role in myopia. Using GFAP as a specific marker, we demonstrated that the morphology of retinal glial cells is altered in FDM mice. We also identified common functional pathways using bioinformatics analyses. The most likely function of glial cells is response to mechanical stimuli. Our findings support the existing hypoxia theory of myopia and provide more evidence for retinal glial function. The future direction of myopia mechanistic research could lead to glial function, because glial cells may constitute an important bridge between neuronal vision information and ocular growth. We propose a potential important role of retinal glia in myopia and call for more detailed research based on our findings.

## Acknowledgements

We are grateful to XinYi Zhao (the First Affiliated Hospital of Zhejiang University) for assistance with guidance for bioinformatics analysis and suggested changes to the manuscript. We thanks to the staffs of core facilities, Zhejiang University school of medicine for assistance with picture taking and analysis.

